# Long-Acting GLP-1 Receptor Agonism Ameliorates Sinoatrial Node Dysfunction by Attenuating Electrophysiological and Structural Remodeling

**DOI:** 10.64898/2026.11.13.766590

**Authors:** Hanfeng Liu, Yangxi Zeng, Rui Zeng, Wei Fan, Can Liu, Zhaoting Li, Bin Liao, Juyi Wan

**Author notes:** These authors contributed equally to the writing of this article (Hanfeng Liu, Yangxi Zeng). Denotes corresponding authors responsible for correspondence and oversight (Bin Liao, Juyi Wan). **Correspondence to:** Juyi Wan, MD, Department of Cardiovascular Surgery, The Affiliated Hospital, Southwest Medical University, Luzhou, China., Bin Liao, MS, Department of Cardiovascular Surgery, The Affiliated Hospital, Southwest Medical University, Luzhou, China.

## Abstract

**Background:** Sinoatrial node dysfunction (SND) causes bradyarrhythmia, while pacemaker therapy does not reverse injury to the sinoatrial node (SAN). Long-acting GLP-1 receptor agonists (GLP-1RAs) raise heart rate, but whether they can repair an injured SAN in SND remains unknown.

**Methods:** In streptozotocin-induced SND rats, we studied an acute phase (SND, GLP-1RA treatment, glycemic control, and GLP-1R antagonist groups) and a chronic phase (SND, GLP-1RA treatment, and glycemic control groups). Endpoints included heart rate, RR interval, heart rate variability, intrinsic heart rate (IHR), corrected sinus node recovery time after adenosine injection (ADSNRT), SAN transcriptomics, histology and immunofluorescence, HL-1 cell oxidative-stress studies, and public single-cell data.

**Results:** At the acute endpoint, the long-acting GLP-1RA efsubaglutide alfa (eSuba) increased heart rate (372.76 ± 11.06 vs 312.64 ± 22.28 beats/min; *P* < 0.0001) and IHR (349.45 ± 10.49 vs 234.79 ± 12.29 beats/min; *P* < 0.0001), and shortened ADSNRT (4.22 ± 2.75 vs 20.68 ± 1.60 s; *P* = 0.0009); improvements in heart rate (349.05 ± 17.73 vs 266.02 ± 13.52 beats/min; *P* < 0.0001) and IHR persisted in the chronic phase. SAN transcriptomics showed reversal of extracellular matrix, inflammatory/immune, redox/mitochondrial, glial, and electrical programs. The GLP-1RA reduced collagen deposition and CCR2a^+^ infiltration, decreased ox-CaMKII/CaMKII, increased SIRT3, restored Ca_v_1.3, Na_v_1.5, SERCA2a and p-RyR2 (Ser2808), and increased p-PKA/PKA. The glycemic control comparator produced limited improvement, and the GLP-1R antagonist attenuated most effects.

**Conclusions:** Long-acting GLP-1R agonism improves SAN function in SND via glucose-independent attenuation of electrophysiological and structural remodeling, an effect that is at least partly GLP-1R dependent.

**Clinical Perspective:** *What is new?:*

- In metabolically stressed rats with established SAN dysfunction, a long-acting GLP-1RA improved SAN function by attenuating electrophysiological and structural remodeling rather than merely raising heart rate, and an insulin comparator matched for glycemic control did not reproduce these benefits—uncoupling SAN repair from glucose lowering.
- Mechanistically, eSuba attenuated SAN oxidative stress and mitochondrial injury (reduced oxidized CaMKII, increased SIRT3) and restored intracellular Ca^2+^ cycling (increased Ca_v_1.3, SERCA2a, and p-RyR2 [Ser2808]), effects that were blunted by GLP-1R blockade.
- SND also reactivated a SAN Schwann/glial transcriptional program (*S100b*; *Ncmap*; etc.) that eSuba reversed, suggesting a modifiable neuro-glial component beyond pacemaker-cell failure.

*What are the clinical implications?:*

- For patients with SND and metabolic disturbances, GLP-1RAs may have therapeutic potential beyond glucose and weight reduction, providing proof of concept for a pharmacological strategy beyond pacemakers.
- The heart-rate effects and safety of GLP-1RAs in patients with bradyarrhythmia warrant reassessment; in the setting of SND, a rise in heart rate may represent a therapeutic effect rather than an adverse reaction.

## Introduction

The sinoatrial node (SAN) is the origin of normal cardiac rhythm. Its function depends on spontaneous electrical activity of pacemaker cells, intercellular coupling, and a microenvironment formed by surrounding fibroblasts, immune cells, blood vessels, and glial components^1, 2^. SND is therefore not only an electrophysiological abnormality of pacemaker cells but also involves alterations in the local tissue architecture and metabolic support that sustain pacemaker function^2, 3^. For patients with an established indication, permanent pacemakers relieve bradycardia-related symptoms, but this is symptomatic therapy and does not reverse the pathological process in the SAN^1^. Strategies targeting SAN structural injury are also being explored: platelet-membrane-engineered hiPSC-sEVs have achieved targeted repair of the fibrotic SAN in preclinical SND models^4^. The molecular mechanisms of SND and precision therapeutic strategies have recently been reviewed^5–7^. Diabetes can induce ion-channel and conduction-system remodeling in the SAN, including reduced HCN channel expression^8^ and SAN microvascular rarefaction^9^, as well as conduction-system remodeling^10–12^. Under metabolic stress, oxidative injury, inflammation, and fibrosis can jointly damage pacemaker cells and their microenvironment; the link between oxidative activation of calcium/calmodulin-dependent protein kinase II (CaMKII) and SAN cell injury and dysfunction has been supported by animal studies^13–15^. These observations suggest that reducing pathological oxidative load and restoring calcium handling in pacemaker cells may be an entry point for improving a damaged SAN.

Glucagon-like peptide-1 receptor agonists (GLP-1RAs) have established cardiovascular benefits^16^ and can increase heart rate^17^. Recent studies have linked this effect to a direct action on the SAN: in pigs and isolated SAN preparations, GLP-1 regulated calcium-clock-related phosphorylation through pacemaker-cell GLP-1R, shortened the pacing cycle, and accelerated diastolic depolarization independently of autonomic tone or the *I_f_* current^18^. However, whether the heart-rate effect of GLP-1RAs is mediated by a direct SAN action or by autonomic reflexes remains debated^19, 20^. Tirzepatide, a dual GIP/GLP-1 receptor agonist, has been shown to modulate cAMP and calcium dynamics in human SAN cells^21^. The expression profile of GLP-1R in the human heart^22^ and its structural pharmacology^23^ have also been described. Importantly, all of these studies examined normal or isolated SAN tissue, and in vivo evidence that GLP-1RAs can improve a SAN with established metabolic injury and structural remodeling is lacking. A recent study showed that semaglutide directly inhibited CaMKII activity, reduced late sodium current, and decreased sarcoplasmic reticulum calcium leak in failing human ventricular myocardium^24^, suggesting a pharmacologically actionable link between GLP-1R and CaMKII; however, that study did not involve the SAN, SIRT3, or structural remodeling.

Efsubaglutide alfa (also known as supaglutide; abbreviated as eSuba here) is a long-acting GLP-1RA composed of two GLP-1 sequences fused to a human IgG2 Fc fragment. Its half-life is approximately 182–215 h, allowing once-weekly dosing^25, 26^. It was approved in China for type 2 diabetes in January 2025^27^ and has been shown to improve hepatic steatosis and fibrosis in a model of metabolic dysfunction-associated steatohepatitis^28^. Given its potent and long-acting GLP-1R agonism, eSuba provides an ideal tool to test whether GLP-1R activation can improve an injured SAN. In the present study, using a streptozotocin (STZ)-induced SND model in Sprague-Dawley rats, we evaluated the functional effects of eSuba in the acute and chronic phases. Using a long-acting insulin active comparator, combined GLP-1R blockade, SAN transcriptomics, pathological and molecular quantification, a cellular oxidative-stress model, and public single-cell data, we tested the hypothesis that eSuba inhibits oxidized CaMKII, upregulates SIRT3, and restores calcium-handling proteins through GLP-1R, thereby improving SAN function and attenuating local structural remodeling.

## Methods

Male Sprague-Dawley rats (8 weeks old, 200–250 g) and male C57BL/6J mice (10 weeks old) were used. In the STZ-induced SND rat model, an acute phase compared four groups (SND, GLP-1RA, glycemic control, and GLP-1R antagonist) and a chronic phase compared three groups (SND, GLP-1RA, and glycemic control); the mouse model was used for 28-day survival and echocardiographic assessment. Endpoints included surface ECG and heart rate variability, intrinsic heart rate after autonomic blockade, corrected sinus node recovery time after adenosine injection (ADSNRT), SAN transcriptomics, histology and immunofluorescence, HL-1 cell oxidative-stress experiments, and public single-cell data. Detailed descriptions of animals and ethics, model induction and dosing, electrophysiological recording, molecular and histological analyses, RNA sequencing and bioinformatics, and single-cell analysis are provided in the Supplemental Material.

### Statistical Analysis

Statistical analyses were performed with GraphPad Prism 10.1.2; the *P* values shown in the figures were calculated and exported by this software. Data are presented as mean ± SD, all tests were two-sided, and *P* < 0.05 was considered statistically significant. Values from different time points were analyzed as independent samples. In the acute phase, heart rate, RR interval, and PNS/SNS indices were compared by two-way ANOVA (group × time), and comparisons at day 0 used the Bonferroni–Šídák correction (four comparisons per family: eSuba versus SND, insulin, and eSuba + exendin 9-39, and eSuba at day 0 versus its own baseline). In the chronic phase, heart rate, RR interval, and PNS/SNS indices at day 28 were compared by one-way ANOVA with Tukey’s multiple-comparison test. IHR was analyzed by one-way ANOVA with Bonferroni–Šídák correction in the acute phase (four groups) or Dunnett’s test in the chronic phase (three groups), and ADSNRT by two-sided unpaired t tests. Survival was analyzed by the Kaplan–Meier method with the log-rank (Mantel–Cox) test. Differential expression used a robust quasi-likelihood GLM in edgeR with a Benjamini– Hochberg FDR < 0.05. Exact group sizes are given in the figure legends, and the tests used for each comparison are specified in the Supplemental Material.

## Results

### eSuba improves SAN function and modulates autonomic balance in acute SND rats

On day 3 after STZ injection (designated d−4), rats were randomized and dosing began; the acute-phase protocol is shown in **Figure 1a**. At 24 h after dosing, in representative recordings, the RR interval change from baseline (ΔRR) was −20 ms in the eSuba group and +26.2 ms in the SND group (**Figure 1b**). At the acute observation endpoint (d0), mean heart rate was significantly higher in the eSuba group than in the SND group (372.76 ± 11.06 vs 312.64 ± 22.28 beats/min, *P* < 0.0001), and the RR interval was shorter (161.07 ± 4.62 vs 192.71 ± 14.06 ms, *P* = 0.0005; **Figure 1c–e**). Heart rate in the insulin glargine group (331.81 ± 19.11 beats/min) was intermediate between the SND and eSuba groups and was significantly lower than in the eSuba group (*P* = 0.0056), whereas heart rate in the eSuba + exendin 9-39 group (313.00 ± 20.46 beats/min) was significantly lower than in the eSuba monotherapy group (*P* < 0.0001; **Figure 1d,e**).

**Figure 1.**
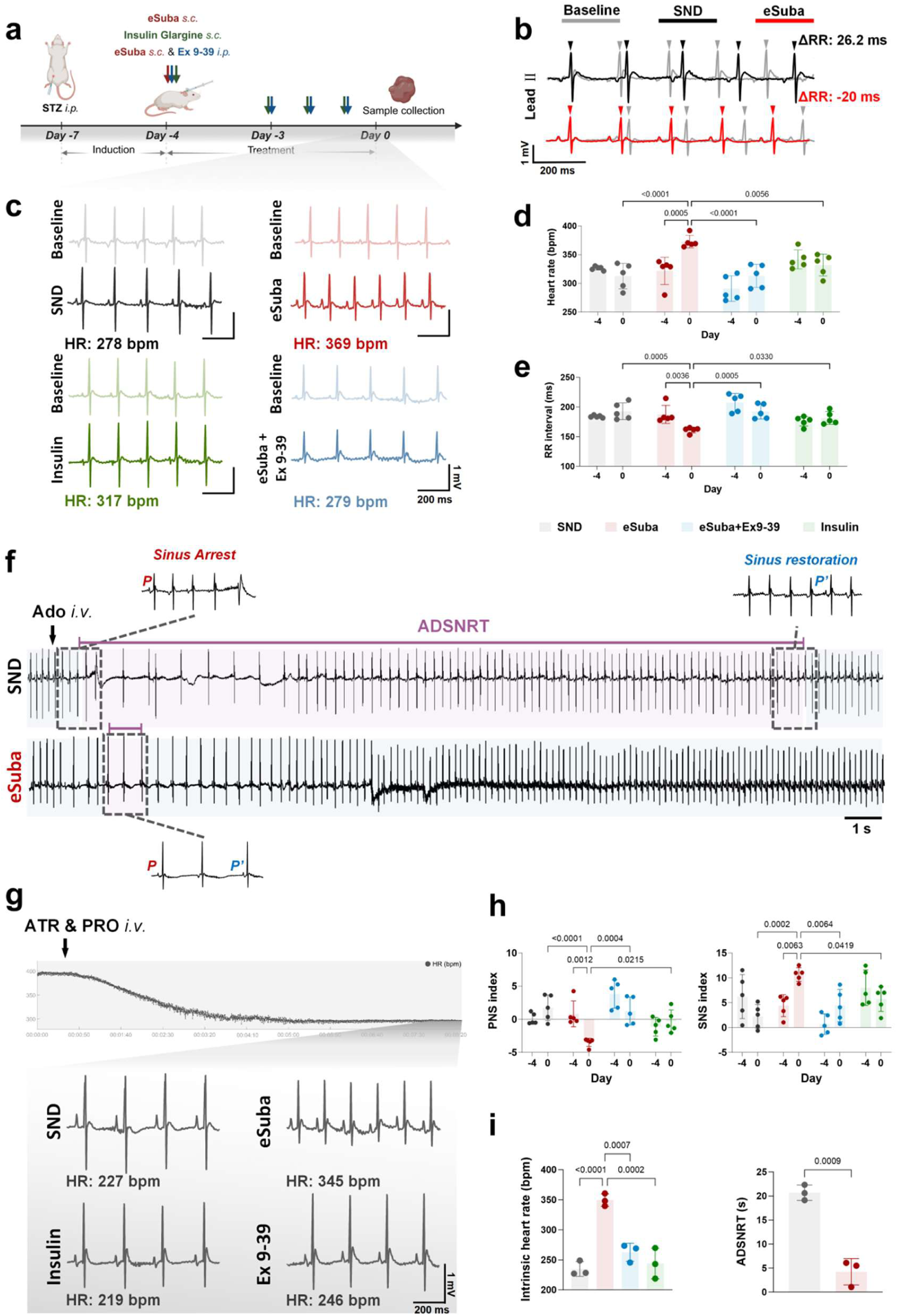
eSuba improves SAN function in acute SND rats. (a) Timeline of STZ injection, dosing, and endpoint tissue collection (STZ d−7; dosing d−4; 24-h recording d−3; endpoint d0).Created in BioRender. (b) Representative lead II ECG recordings before and 24 h after dosing; ΔRR = RR interval at 24 h after dosing − baseline RR interval in the same animal (SND +26.2 ms; eSuba −20 ms); scale bars are indicated in the figure. (c) Representative ECGs of each group at baseline and at the acute observation endpoint. (d,e) Mean heart rate (d) and RR interval (e) at d−4 and d0. (f) Representative ECG recordings showing ADSNRT in the eSuba and SND groups at the acute endpoint (loss and recovery of sinus P waves). (g) Representative ECG recordings of IHR after dual blockade with atropine and propranolol at the acute endpoint. (h) PNS index and SNS index at the observation endpoint. (i) Quantitative comparison of IHR and ADSNRT at the observation endpoint. n = 5 per group for heart rate/RR/HRV and n = 3 per group for IHR/ADSNRT; data are mean ± SD; (d,e,h) Two-way ANOVA (group × time) with Bonferroni–Šídák-adjusted comparisons; (i) one-way ANOVA with Bonferroni–Šídák correction (IHR) and a two-sided unpaired t test (ADSNRT); specific *P* values are shown in the figure.

Analysis of autonomic indices showed that the PNS index was lower in the eSuba group than in the SND group (−3.49 ± 0.62 vs 1.74 ± 1.95, *P* < 0.0001), and the SNS index was higher (10.68 ± 1.39 vs 2.21 ± 2.41, *P* = 0.0002), indicating a shift of autonomic balance toward sympathetic predominance (**Figure 1h**).

After dual autonomic blockade, IHR was significantly higher in the eSuba group than in the SND group (349.45 ± 10.49 vs 234.79 ± 12.29 beats/min, *P* < 0.0001; **Figure 1g,i**), and was also higher than in the eSuba + exendin 9-39 group (262.68 ± 15.04 beats/min, *P* = 0.0007) and the insulin group (243.93 ± 25.45 beats/min, *P* = 0.0002); IHR in the insulin group remained close to the SND level (**Figure 1i**). After adenosine injection, ADSNRT was significantly shorter in the eSuba group than in the SND group (4.22 ± 2.75 vs 20.68 ± 1.60 s, *P* = 0.0009; **Figure 1f,i**). These results indicate that eSuba improved pacemaker function already in the acute phase, and that insulin with comparable glucose lowering did not reproduce most of these effects; the time-domain distribution and Poincaré analysis of HRV are shown in **Figure S1**.

### eSuba persistently improves SAN function and remodels autonomic balance in the chronic phase

The chronic-phase protocol is shown in **Figure 2i**. Serial observations showed that heart rate in the eSuba group increased from d7 and was maintained through d28, whereas heart rate in the SND group progressively declined and the RR interval changed in the opposite direction (**Figure 2a**); representative baseline and endpoint ECGs are shown in **Figure 2c**. At d28, mean heart rate (349.05 ± 17.73 vs 266.02 ± 13.52 beats/min, *P* < 0.0001; **Figure 2g**), IHR (328.36 ± 2.39 vs 255.82 ± 5.14 beats/min, *P* = 0.0137; **Figure 2f,h**), and ADSNRT (6.31 ± 1.82 vs 21.13 ± 5.79 s, *P* = 0.0134; **Figure 2d,h**) in the eSuba group differed significantly from those in the SND group; IHR was also higher than in the insulin group (217.50 ± 38.71 beats/min, *P* = 0.0017; **Figure 2h**). At d28, the PNS index was lower in the eSuba group than in the SND group (−1.68 ± 1.29 vs 6.80 ± 2.07, *P* < 0.0001), and the SNS index was higher (8.11 ± 2.90 vs −2.55 ± 1.57, *P* < 0.0001; **Figure 2b,g**). In the insulin group, heart rate increased only transiently at d7 and did not differ significantly from the SND group at d28 (286.67 ± 8.89 beats/min, *P* = 0.0859; **Figure 2a,g**). Histograms of RR interval distributions and Poincaré analyses for each group are shown in **Figure 2e**.

**Figure 2.**
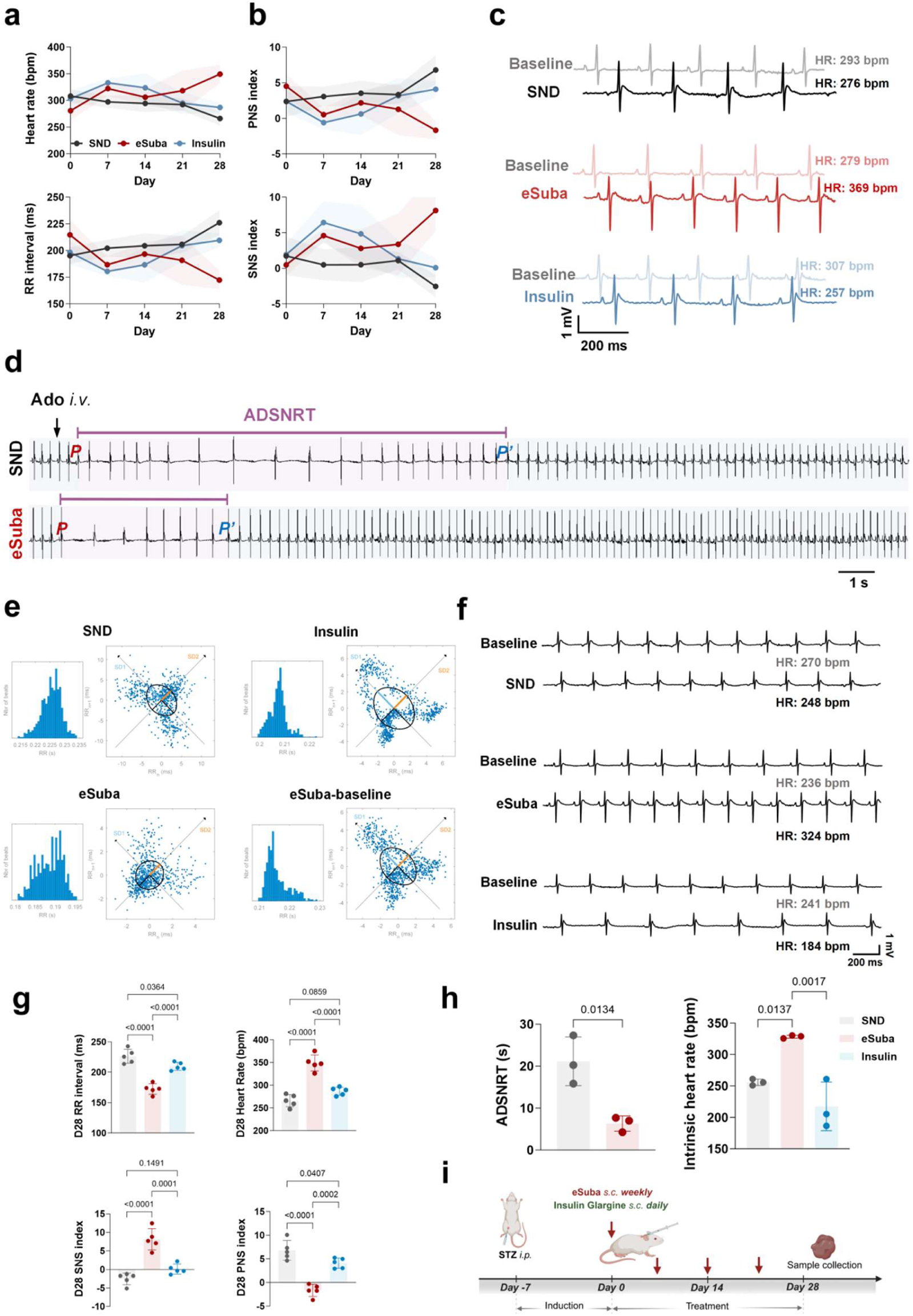
eSuba persistently improves SAN function in the chronic phase. (a) Changes in heart rate and RR interval at d0, d7, d14, d21, and d28 (SND, eSuba, and insulin groups). (b) Changes in PNS index and SNS index over the same period. (c) Representative ECGs of each group at baseline and at the observation endpoint. (d) Representative ECG recordings showing ADSNRT in the eSuba and SND groups at the observation endpoint. (e) Time-domain HRV histograms and Poincaré plots for the SND, insulin, eSuba, and eSuba-baseline groups. (f) Representative ECG recordings of IHR in each group at the observation endpoint. (g) Statistical plots of heart rate, RR interval, and PNS/SNS indices at the observation endpoint. (h) Statistical plots of IHR and ADSNRT at the observation endpoint. (i) Schematic of the chronic-phase experimental protocol. Created in BioRender. n = 5 per group for heart rate/RR/HRV and n = 3 per group for IHR/ADSNRT; data are mean ± SD; (g) One-way ANOVA with Tukey’s multiple-comparison test; (h) one-way ANOVA with Dunnett’s test (IHR) and a two-sided unpaired t test (ADSNRT); specific *P* values are shown in the figure.

### SAN transcriptomics reveals SND-associated multi-module remodeling programs

RNA-seq was performed on SAN tissue from the SND and eSuba groups at the chronic endpoint (four biological replicates per group; the heatmap and PCA show three representative replicates per group). Across the four replicates per group, 277 genes were differentially expressed (FDR < 0.05 and |log2FC| ≥ 1; 47 upregulated and 230 downregulated). **Figure S14a** shows a representative heatmap of the differentially expressed genes (7 upregulated and 43 downregulated in the displayed subset). The volcano plot is shown in **Figure S14c**. Principal component analysis of 12,117 genes in six representative samples (three per group) showed that PC1, PC2, and PC3 explained 49.22%, 17.14%, and 12.81% of the variance, respectively (**Figure S14b**). Gene Ontology enrichment of the differentially expressed genes was dominated by extracellular matrix/structural terms (extracellular matrix, collagen-containing extracellular matrix, extracellular region and space, and cell periphery), lipid and organic acid metabolic processes, and receptor–ligand/immunoglobulin binding (**Figure S14d**). GSEA showed enrichment of extracellular matrix, superoxide anion generation, and immune effector gene sets in SND (NES −2.33 to −1.80; FDR ≤ 0.022), with glial activation at FDR = 0.060; in eSuba, mitochondrial membrane permeability, ATP biosynthetic process, and proton-motive-force-driven ATP synthesis were enriched (NES 1.53–1.64; FDR 0.18–0.23), whereas glucose metabolism, DNA repair, and mitochondrial matrix were nominally enriched (FDR 0.27–0.48) (**Figure S14e**). KEGG enrichment included ECM–receptor interaction, focal adhesion, and PI3K–Akt signaling; Reactome enrichment was dominated by extracellular matrix organization and collagen formation (**Figure S3**). In a protein–protein interaction network constructed from 146 candidate genes potentially involved in SND pathogenesis (543 edges), high-connectivity nodes included *Spp1*, *Fbn1*, *Loxl1*, *Ccl2*, and *Cx3cr1* (**Figure S14f**). Metabolism-related candidate genes (e.g., increased *Slc2a4* and *Ppara*; decreased *Ldlr* and *Fabp4*) are shown in **Figure S7**, suggesting a shift in substrate utilization. The following sections use this transcriptomic candidate set as a starting point for histological and molecular validation of each module.

### eSuba reverses extracellular matrix/fibrosis-related transcriptional programs and reduces SAN collagen deposition

Transcriptomics showed that extracellular matrix-related genes were globally downregulated in the eSuba group: collagens (*Col1a1* and *Col3a1*), elastin (*Eln*), fibrillin (*Fbn1*), proteoglycans (*Fmod*, *Lum*, and *Aspn*), and genes involved in collagen maturation/crosslinking (*Serpinh1*, *Pcolce*, *Loxl1/2*, and *Mfap5*) showed reduced expression (**Figure 3a**; extracellular matrix GSEA is shown in **Figure S8a,b**). Histological validation showed that the proportion of Masson-positive area in the SAN was significantly lower in the eSuba group than in the SND group (19.46 ± 2.03% vs 49.35 ± 4.90%, *P* < 0.0001), and the proportion of Sirius red-positive area was also significantly lower (21.15 ± 7.46% vs 51.20 ± 4.74%, *P* < 0.0001; **Figure 3c,d**). Compared with the insulin group, the Masson- and Sirius red-positive areas were further reduced in the eSuba group (*P* = 0.0004 and *P* = 0.0005, respectively; **Figure 3d**). The corresponding values in the insulin group were 40.51 ± 9.27% and 44.11 ± 7.67%, which did not differ significantly from those in the SND group (*P* = 0.10 and *P* = 0.26; **Figure 3d**). Mean fluorescence intensities of vimentin (1.56 ± 0.61 vs 11.43 ± 2.59) and COL3A1 (2.28 ± 1.42 vs 20.38 ± 3.97) in the eSuba group were also significantly lower than in the SND group (both *P* < 0.0001; **Figure 3e,f**). Vimentin and COL3A1 were also lower in the insulin group than in the SND group, but the reductions were smaller than those in the eSuba group (**Figure 3e,f**). In addition, Masson and Sirius red staining of ventricular myocardium at the chronic endpoint showed increased interstitial fibrosis in the SND group that was markedly attenuated in the eSuba group (**Figure S8c**), suggesting that the antifibrotic effect of eSuba is not limited to the SAN. Co-staining for Cx43 and CaMKIIδ was used to define the boundary between SAN and atrial tissue (**Figure 3b**), providing an anatomical basis for regional quantification.

**Figure 3.**
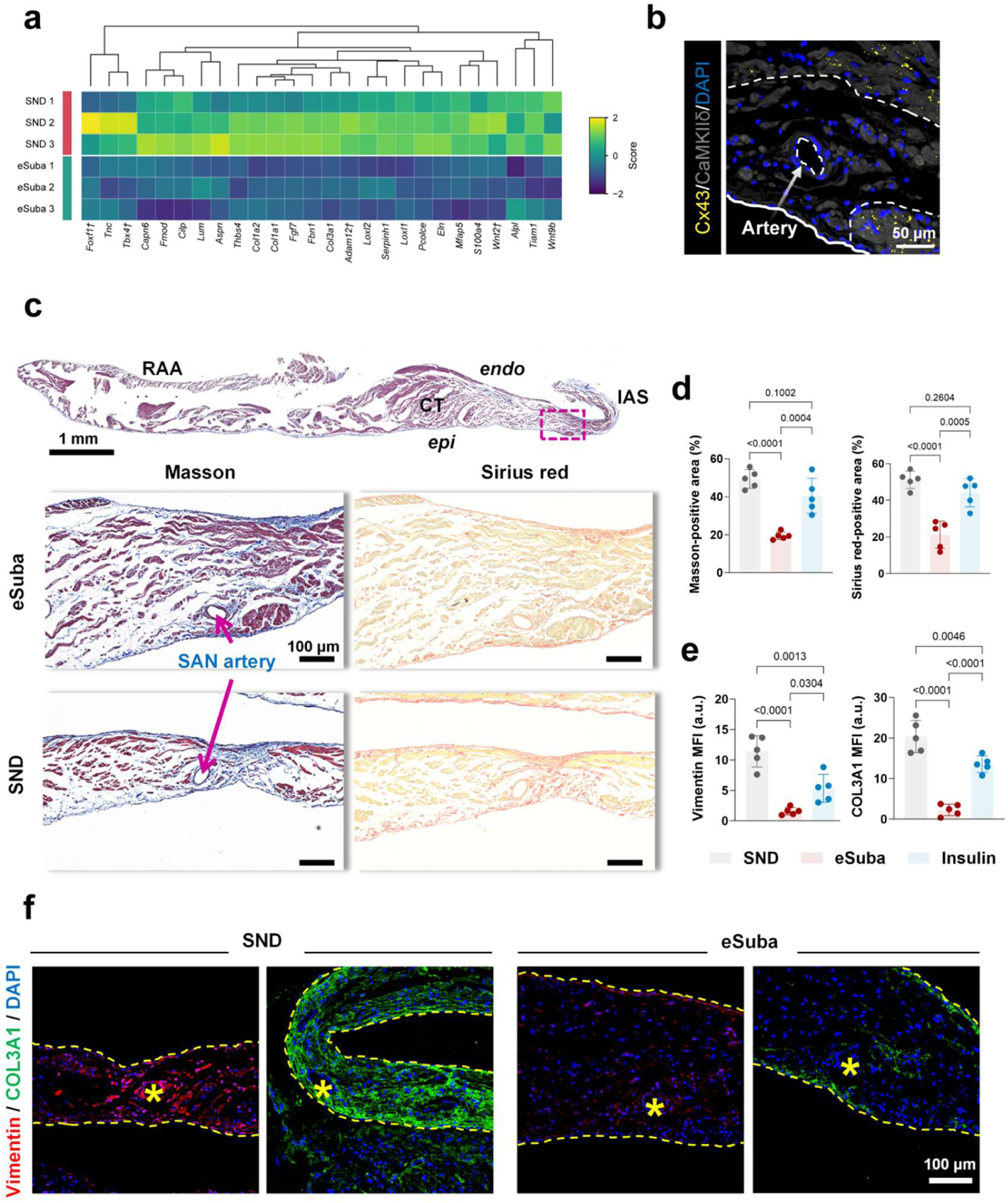
eSuba attenuates SAN fibrosis in chronic SND rats. (a) Heatmap of fibrosis/extracellular matrix-related genes among significantly differentially expressed genes (three representative replicates per group are shown). (b) Co-staining for Cx43 (yellow) and CaMKIIδ (gray) used to define the boundary between SAN and atrial tissue; the area outlined by the white dashed line indicates the Cx43-positive SAN region, and arrows indicate the SAN artery (scale bar, 50 μm). (c) Masson and Sirius red staining of the SAN region in the SND and eSuba groups (RAA, right atrial appendage; CT, crista terminalis; IAS, interatrial septum; arrows indicate the SAN artery; low-magnification scale bar, 1 mm; high-magnification scale bar, 100 μm). (d) Quantification of Masson- and Sirius red-positive area (n = 5 per group). (e) Quantification of mean fluorescence intensity of vimentin and COL3A1 (n = 5 per group). (f) Representative immunofluorescence images of vimentin (red) and COL3A1 (green) (scale bar, 100 μm); the yellow dashed line outlines the SAN region and yellow stars indicate the SAN artery. Data are mean ± SD; between-group comparisons were made by one-way ANOVA with Tukey’s multiple-comparison test; specific *P* values are shown in the figure.

### eSuba reverses inflammation/immune-related transcriptional programs and reduces inflammatory cell infiltration in the SAN

Transcriptomics showed that inflammation- and immune-effector-related genes were significantly downregulated in the eSuba group, including chemokines and their receptors (e.g., *Ccl2*, *Ccl6*, and *Cx3cr1*), pro-inflammatory mediators and tissue-remodeling factors (e.g., *Il1a* and *Spp1*), and immune-cell markers (e.g., *Ly6c* and *Cd209f*) (**Figure S15a**); Gene Ontology immune-effector and inflammatory-response gene sets were enriched in the SND group (**Figure S15b**; **Figure S5c**). Histological validation showed that in the acute phase, the proportions of CCR2a^+^ cells (0.70 ± 0.65% vs 25.00 ± 5.24%, *P* < 0.0001) and CD68^+^ cells (3.05 ± 2.02% vs 21.70 ± 6.39%, *P* < 0.0001) in the SAN were significantly lower in the eSuba group than in the SND group (**Figure S15c,d**). In the chronic phase, the proportion of CCR2a^+^ cells was also significantly reduced (1.48 ± 1.41% vs 16.96 ± 2.94%, *P* = 0.0012; **Figure S15e,f**), and CD68^+^ cells showed a decreasing trend (6.20 ± 1.76% vs 12.93 ± 3.16%, *P* = 0.060; **Figure S15e,f**). The proportions of CCR2a^+^ and CD68^+^ cells in the eSuba + exendin 9-39 group were significantly higher than in the eSuba monotherapy group (CCR2a^+^ *P* < 0.0001; CD68^+^ *P* = 0.027; **Figure S15d**), whereas the insulin group was close to the SND group. Plasma TNF-α was higher in the SND group than in the baseline group (*P* = 0.0335), was lower in the eSuba group than in the SND group (*P* = 0.0444) and remained close to baseline, and did not differ between the SND and insulin groups (*P* = 0.5452); IL-6 did not differ among groups (**Figure S5b**). H&E staining showed myocardial inflammatory foci and CD68^+^ cell infiltration that were markedly reduced by eSuba and partially reversed by exendin 9-39 (**Figure S4a,b**), and multiorgan inflammatory injury in the SND group (lung, liver, kidney, and spleen) was attenuated by eSuba (**Figure S5a**).

### eSuba reverses oxidative-stress/mitochondria-related transcriptional programs and inhibits ox-CaMKII

Transcriptomics showed favorable changes in genes related to mitochondrial biogenesis and antioxidant defense in the eSuba group: *Ppargc1a* and *Nr4a1* were upregulated, whereas *Nnmt*, *Ucp3*, *Ncf4*, *Duox1*, and *Atf5* were downregulated (**Figure 4a**); Gene Ontology gene sets related to regulation of mitochondrial membrane permeability and superoxide anion generation were enriched in the SND group (**Figure 4b**). Histological validation showed that in the acute phase, the mean fluorescence intensity of ox-CaMKII in the SAN was reduced by approximately 88% in the eSuba group relative to the SND group (7.85 ± 3.88 vs 66.69 ± 6.09 a.u., *P* < 0.0001), the ox-CaMKII/CaMKII ratio was reduced by approximately 95% (0.15 ± 0.07 vs 3.02 ± 1.12, *P* < 0.0001), and total CaMKII signal was increased (52.45 ± 9.86 vs 24.09 ± 6.92; **Figure 4c,d**). Mean SIRT3 fluorescence intensity was significantly increased (44.11 ± 15.33 vs 0.16 ± 0.17 a.u., *P* < 0.0001), and 8-OHdG was significantly reduced (0.53 ± 0.14 vs 20.66 ± 2.67 a.u., *P* < 0.0001; **Figure 4c,d**). After co-administration of exendin 9-39, the ox-CaMKII/CaMKII ratio rose to 2.00 ± 0.49 (*P* = 0.0017 vs eSuba), SIRT3 decreased (*P* < 0.0001 vs eSuba), and 8-OHdG increased (*P* < 0.0001 vs eSuba; **Figure 4d**); in the insulin group, the ratio remained elevated (3.06 ± 0.41; *P* < 0.0001 vs eSuba), SIRT3 remained low (*P* < 0.0001 vs eSuba), and 8-OHdG was higher than in the eSuba group (9.34 ± 3.22; *P* = 0.036; **Figure 4d**). In the chronic phase, the direction of changes was consistent: ox-CaMKII decreased by approximately 77%, the ratio decreased by approximately 85%, SIRT3 increased from 2.06 ± 0.80 to 33.78 ± 9.60 a.u., and 8-OHdG decreased by approximately 92% (ratio *P* = 0.0014; SIRT3 *P* < 0.0001; **Figure 4e,f**). The mechanism schematic and HL-1 cell immunoblotting are shown in **Figure 4h** and **Figure 4g,i**, respectively: in the eSuba group, the ox-CaMKII/CaMKII ratio and relative expression of dynamin-related protein 1 (DRP1) were lower than in the H_2_O_2_ model group, and SIRT3 was higher than in the model group; co-treatment with exendin 9-39 partially reversed these trends. Because each group contained only three independent culture batches, these comparisons are exploratory: the ox-CaMKII/CaMKII difference was not statistically significant (*P* = 0.081), and only the DRP1 difference remained significant after correction (*P* < 0.001). These data are therefore reported as directionally supportive evidence consistent with the tissue results. The mechanism schematic (**Figure 4h**) summarizes that eSuba, acting through GLP-1R, inhibits ROS/ox-CaMKII and Ca^2+^ overload, upregulates SIRT3, suppresses DRP1-related mitochondrial fission, and modulates transcriptional programs such as *Nr4a1*/*Ppargc1a*, thereby reducing oxidative DNA damage and pacemaker-cell apoptosis.

**Figure 4.**
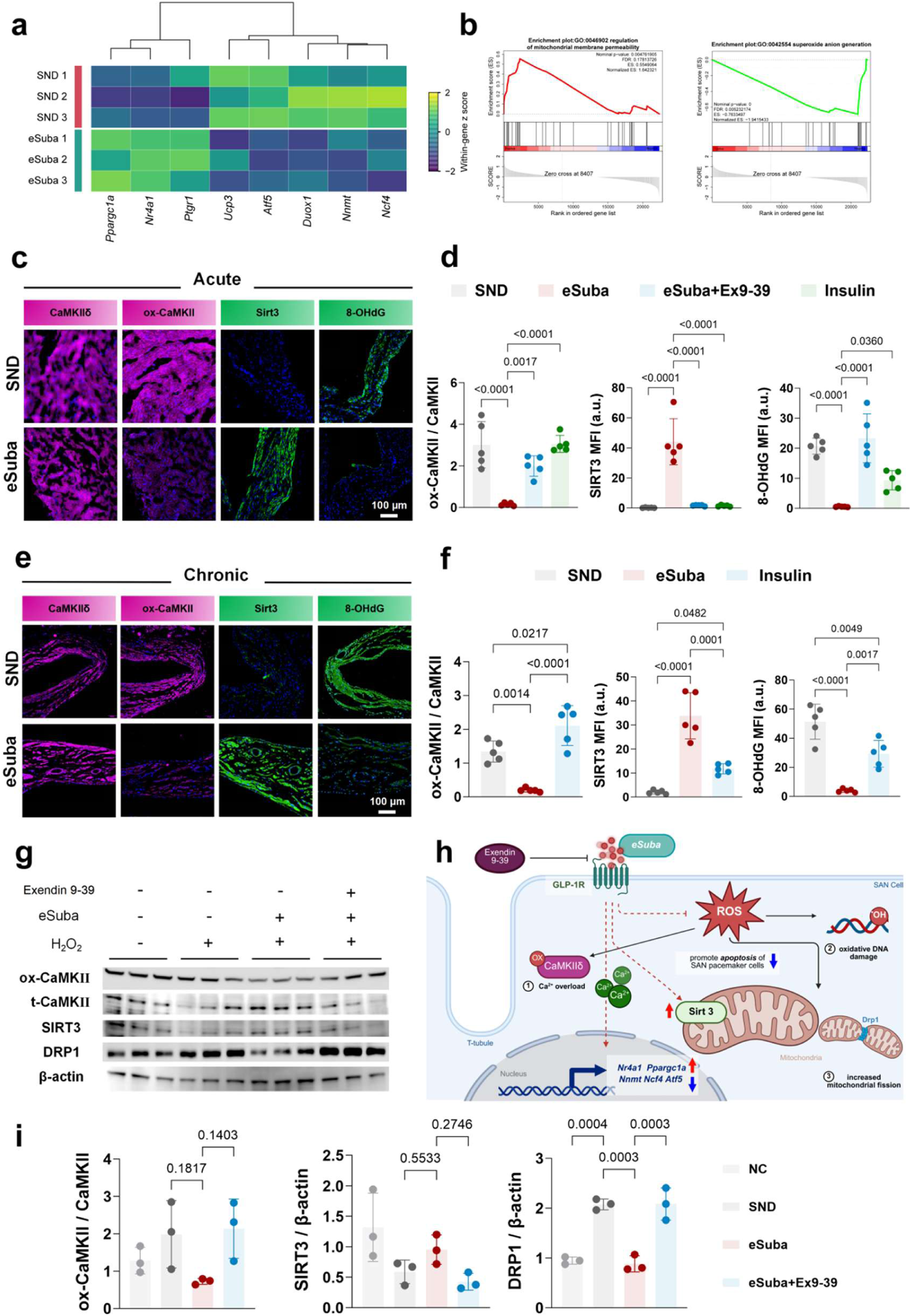
eSuba reduces oxidative stress injury in the SAN and modulates mitochondria-related molecules. (a) Heatmap of differentially expressed genes related to mitochondrial function/metabolism and oxidative stress (three representative replicates per group are shown). (b) GSEA plots for Gene Ontology gene sets. (c) Representative immunofluorescence images of CaMKIIδ, ox-CaMKII, SIRT3, and 8-OHdG in the SAN during the acute phase (scale bar, 100 μm). (d) Quantification of these indices during the acute phase (n = 5 per group). (e) Corresponding representative immunofluorescence images during the chronic phase (scale bar, 100 μm). (f) Quantification during the chronic phase (n = 5 per group). (g) Representative immunoblots of ox-CaMKII, total CaMKII, SIRT3, DRP1, and β-actin in an oxidative-stress-induced HL-1 cell model. (h) Mechanism schematic: eSuba, through GLP-1R, inhibits ROS/ox-CaMKII and Ca^2+^ overload, promotes SIRT3 expression, suppresses DRP1-mediated mitochondrial fission, and modulates nuclear gene transcription such as *Nr4a1*/*Ppargc1a*, thereby reducing oxidative DNA damage and pacemaker-cell apoptosis. Created in BioRender. (i) Normalized quantification of immunoblots (n = 3 independent culture batches per group). (d,f) One-way ANOVA with Tukey’s multiple-comparison test; (i) one-way ANOVA with Holm–Šídák adjustment (the ox-CaMKII/CaMKII ANOVA was not significant, *P* = 0.081; comparisons are exploratory). Data are mean ± SD; specific *P* values are shown in the figure.

### eSuba reverses electrical-activity/calcium-handling transcriptional programs and restores ion-channel protein expression

Transcriptomics showed favorable changes in genes related to electrical activity and calcium handling in the eSuba group: *Cacna1d* (Ca_v_1.3), *Kcna2*, *Kcnip4*, *Pln*, *Myzap*, *Grid1*, and *Gria3* were upregulated, and *Hcn1*, *Hcn2*, *Hcn4*, *Ryr2*, and *Ryr3* tended to be upregulated (**Figure 5a**; mechanism schematic in **Figure 5b**). The schematic in **Figure 5b** shows that eSuba, through GLP-1R, promotes the expression of Ca_v_1.3, Na_v_1.5, and SERCA2a and the phosphorylation of RyR2 at Ser2808, and upregulates nuclear genes including *Cacna1d*, *Fgf12*, *Kcnip4*, and *Myzap*, coordinating intracellular Ca^2+^ cycling with membrane electrical activity; exendin 9-39 blocks this process. Histological validation showed that in the acute phase, SAN SERCA2a (60.03 ± 4.42 vs 25.50 ± 6.16, *P* < 0.0001), Ca_v_1.3 (22.34 ± 5.10 vs 5.85 ± 1.83, *P* < 0.0001), and the p-RyR2 (Ser2808)/RyR2 ratio (0.52 ± 0.08 vs 0.03 ± 0.01, *P* < 0.0001) were significantly higher in the eSuba group than in the SND group (**Figure 5c,e**); Nav1.5 (*Scn5a*) was significantly higher in the eSuba group than in the SND group (17.80 ± 5.84 vs 8.12 ± 2.28, *P* = 0.0011) and than in the exendin 9-39 and insulin groups (both *P* < 0.001; **Figure 5c,e**). In the chronic phase, SERCA2a (38.94 ± 5.78 vs 7.56 ± 2.59, *P* < 0.0001), Ca_v_1.3 (23.51 ± 6.72 vs 11.78 ± 1.56, *P* = 0.0019), Na_v_1.5 (20.68 ± 4.76 vs 6.45 ± 1.78, *P* < 0.0001), and the p-RyR2/RyR2 ratio (2.04 ± 0.26 vs 0.15 ± 0.09, *P* < 0.0001) were all significantly higher in the eSuba group than in the SND group (**Figure 5c,f**). After co-administration of exendin 9-39, these proteins and ratios were reduced by approximately 65%–88% relative to the eSuba group (all *P* < 0.01; **Figure 5e,f**). Immunoblotting and quantification of SERCA2a and Ca_v_1.3 in HL-1 cells are shown in **Figure 5d,g**; relative expression was higher in the eSuba group than in the H_2_O_2_ model group and decreased after exendin 9-39 co-treatment; however, each group contained only three independent culture batches, so these data are reported as exploratory supportive evidence. In addition, Cx43 density was lower and Cx43 was redistributed from intercalated discs to the lateral cardiomyocyte membrane in the SND group, and eSuba partially reversed these changes (**Figure S9**), suggesting improved intercellular coupling. An increased p-RyR2 (Ser2808) ratio suggests enhanced PKA-dependent phosphorylation of RyR2. Consistent with this, cAMP-synthesis genes (*Adrb1*, *Adcy5*, *Adcy6*, and *Popdc2*) were upregulated and phosphodiesterase genes (*Pde5a*, *Pde3b*, *Pde1b*, and *Adra2a*) were downregulated in the eSuba group (**Figure 6a,b**), suggesting increased cAMP synthesis and reduced degradation. Further protein validation showed that at the chronic endpoint, the SAN p-PKA/PKA ratio was higher in the eSuba group than in the SND and Sham groups (3.80 ± 0.71 vs 0.77 ± 0.10 and 0.83 ± 0.25; both *P* < 0.0001; **Figure 6c,d**); the ratio was also increased in the insulin group (2.93 ± 0.06) but was lower than in the eSuba group, with the difference not reaching statistical significance (*P* = 0.0581; **Figure 6c,d**). These findings support that eSuba promotes RyR2 Ser2808 phosphorylation through the canonical GLP-1R–cAMP–PKA axis, thereby improving SAN calcium cycling.

**Figure 5.**
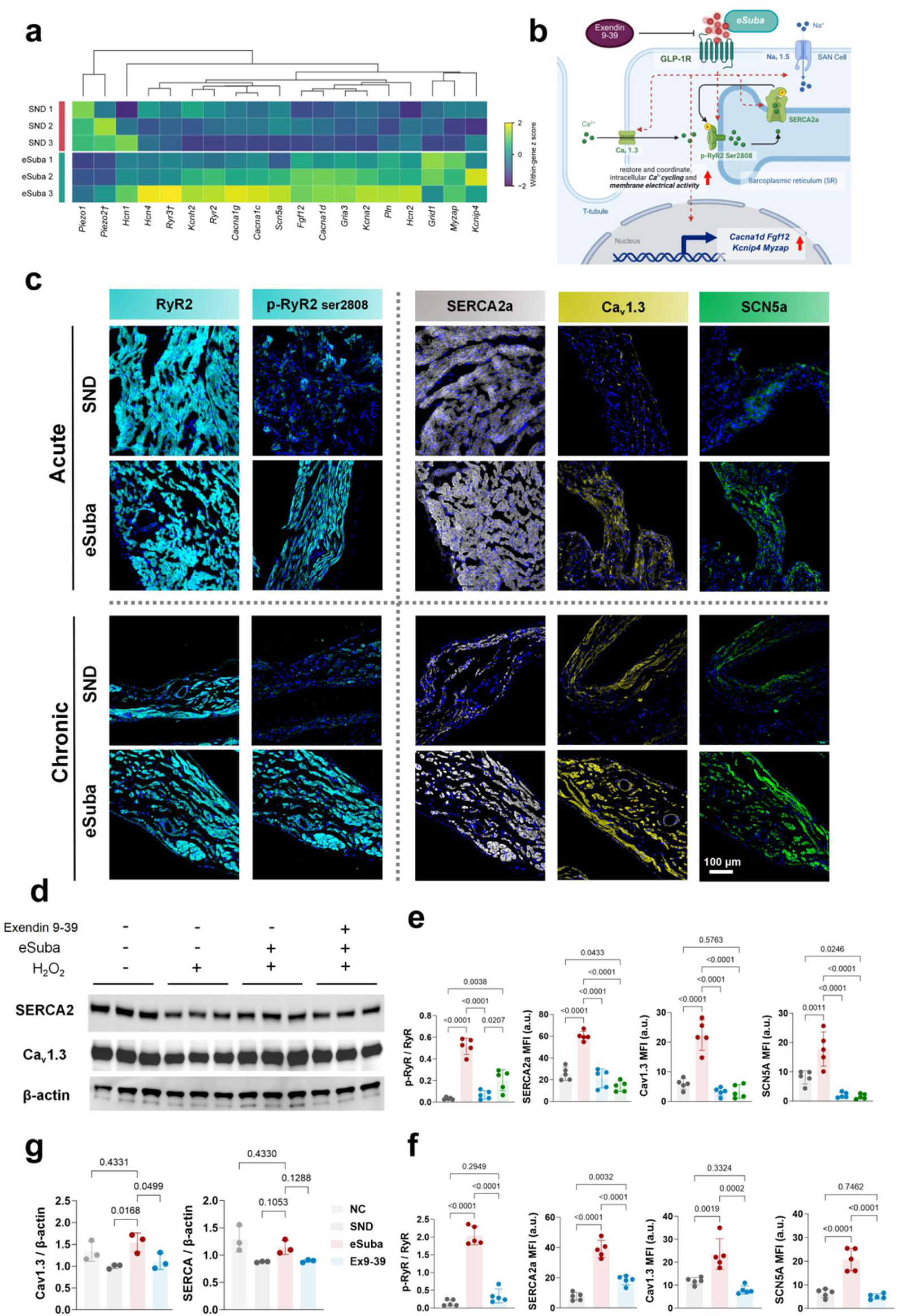
eSuba restores SAN calcium-handling proteins and PKA phosphorylation. (a) Heatmap of differentially expressed genes related to SAN automaticity and electrical conduction (three representative replicates per group are shown). (b) Mechanism schematic: eSuba, through GLP-1R, promotes the expression of Ca_v_1.3, Na_v_1.5, and SERCA2a and the phosphorylation of RyR2 at Ser2808, and upregulates nuclear genes including *Cacna1d*, *Fgf12*, *Kcnip4*, and *Myzap*, coordinating intracellular Ca^2+^ cycling with membrane electrical activity; exendin 9-39 can block this process. Created in BioRender. (c) Representative immunofluorescence images of RyR2, p-RyR2 (Ser2808), SERCA2a, Ca_v_1.3, and Na_v_1.5 (*Scn5a*) in the SAN at the acute and chronic observation endpoints (scale bar, 100 μm). (d) Representative immunoblots of SERCA2a and Ca_v_1.3 in HL-1 cells. (e) Quantification of immunofluorescence during the acute phase (n = 5 per group). (f) Quantification of immunofluorescence during the chronic phase (n = 5 per group). (g) Statistical plots of immunoblotting in HL-1 cells (n = 3 independent culture batches per group). (e,f) One-way ANOVA with Tukey’s multiple-comparison test; (g) one-way ANOVA with Dunnett’s test (eSuba as the control). Data are mean ± SD; specific *P* values are shown in the figure.

**Figure 6.**
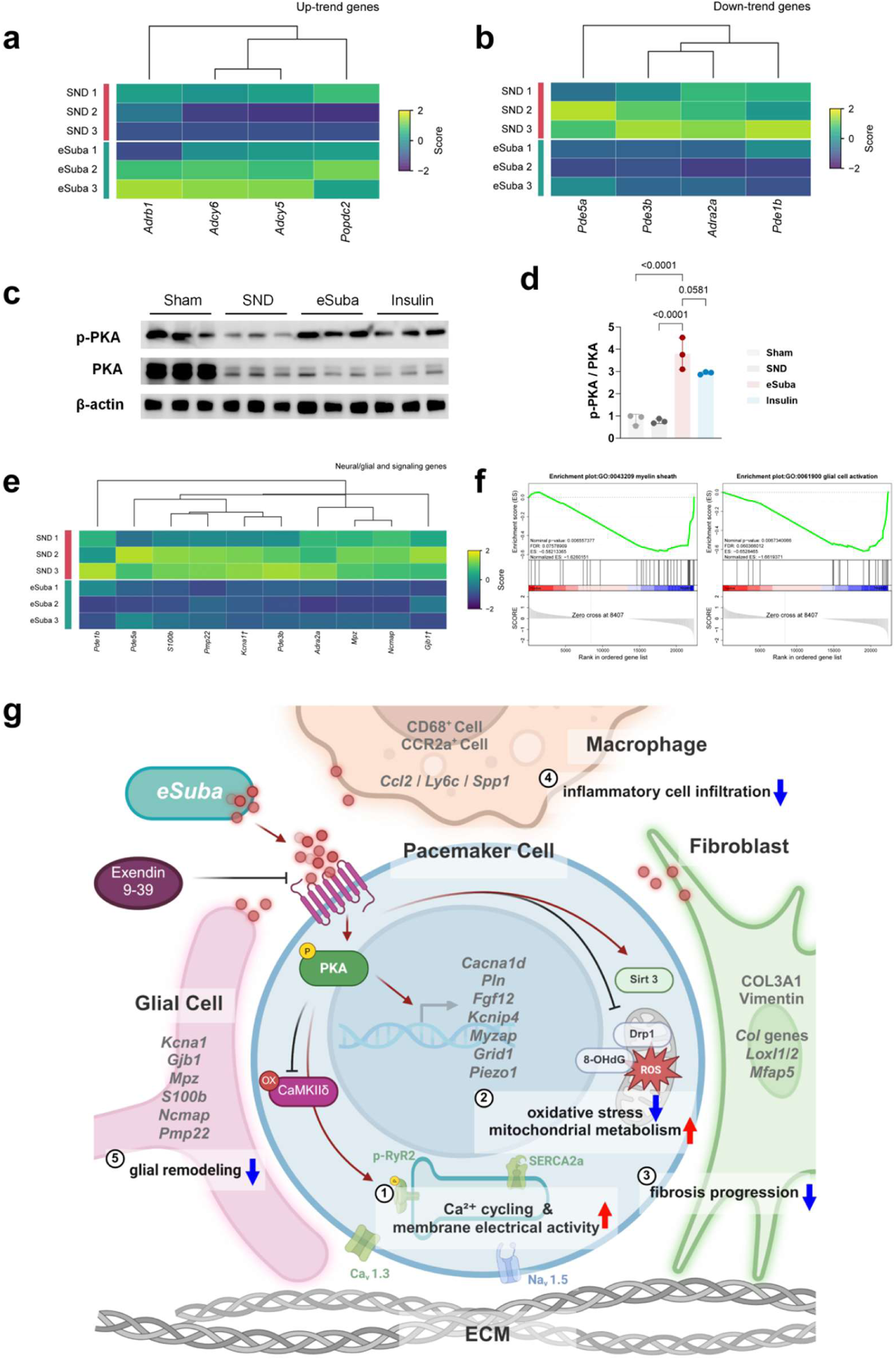
Effects of eSuba on the cAMP-PKA pathway, glia-related transcription, and the integrated working model. (a) Heatmap of upregulated genes related to the cAMP-PKA pathway (*Adrb1*, *Adcy5*, *Adcy6*, and *Popdc2*) (three representative replicates per group are shown). (b) Heatmap of downregulated genes (*Pde5a*, *Pde3b*, *Pde1b*, and *Adra2a*) (three representative replicates per group are shown). (c) Immunoblots of p-PKA, total PKA, and β-actin in SAN tissue at the chronic observation endpoint (Sham, SND, eSuba, and insulin groups; n = 3 per group). (d) Normalized quantification of p-PKA/PKA; one-way ANOVA with Dunnett’s test (eSuba as the control). (e) Heatmap of glia/myelin-related genes (three representative replicates per group are shown). (f) GSEA plot of the related Gene Ontology gene set. (g) Integrated summary model: direct effects of eSuba on SAN pacemaker cells, together with improvement of the surrounding microenvironment, promote recovery of pacemaker function. Created in BioRender. Data are mean ± SD; statistical methods are described in the Supplemental Material; specific *P* values are shown in the figure.

### eSuba reverses glial/myelin-related transcriptional programs

In the transcriptome, Schwann cell/myelin- and glial activation-related transcripts were significantly downregulated in the eSuba group, including *Mpz*, *Pmp22*, *Ncmap*, *Gjb1*, *S100b*, and *Kcna1* (**Figure 6e**); GSEA showed enrichment of myelin- and glial activation-related gene sets in the SND group (**Figure 6f**). Public single-cell data from the developing mouse SAN showed the presence of Schwann/glial lineage cells in the SAN and adjacent atrial region (**Figure S11**), providing a cellular context for these transcriptional changes. These results suggest that SND is accompanied by local glial activation/myelin remodeling in the SAN, which is attenuated by eSuba; this part is a transcriptome-level finding and requires direct protein and cellular validation. Integrating the functional, molecular, and pharmacological blockade results, we propose a working model of how eSuba improves SAN function (**Figure 6g**).

### eSuba improves cardiac structure and function

In the STZ-induced mouse SND model, all 5 animals in the SND group died during the 28-day observation period, whereas 4 of 5 animals in the eSuba group survived (**Figure S2a**; *P* = 0.0263); heart-rate changes in mice are shown in **Figure S2b**. Supplemental echocardiography showed that left ventricular ejection fraction (EF; 78.97 ± 2.25% vs 50.91 ± 2.48%, *P* < 0.0001), fractional shortening (FS; 44.86 ± 1.72% vs 24.72 ± 2.03%, *P* < 0.0001), and ratio of early-to-late mitral inflow velocities (E/A; 1.446 ± 0.053 vs 1.016 ± 0.075, *P* < 0.0001) were higher in the eSuba group than in the SND group (n = 5 per group; **Figure S12a,b**); wheat germ agglutinin (WGA) staining showed that cardiomyocyte cross-sectional area was smaller in the eSuba group than in the SND group (177.04 ± 18.37 vs 735.20 ± 44.43 μm2/cell, *P* < 0.0001; **Figure S12c**). Blood glucose monitoring showed that both the eSuba and insulin groups maintained glucose at similar normal levels (approximately 7–8 mmol/L) during the observation period, whereas the SND group remained hyperglycemic (**Figure S13**), providing a key basis for comparing efficacy under comparable glucose-lowering conditions. Serum ALT and BUN were higher in the SND group than in the baseline group (*P* = 0.0073 and *P* = 0.0014, respectively); ALT and BUN did not differ significantly between the SND group and either the eSuba or insulin group, and AST and Cr did not differ significantly among groups (**Figure S6**).

## Discussion

eSuba improved SAN function in both the acute and chronic phases and reversed the extracellular matrix, inflammatory/immune, redox/mitochondrial, electrical/calcium-handling, and glial programs, with reduced fibrosis and inflammatory infiltration, lower ox-CaMKII, higher SIRT3, restored calcium-handling proteins, and enhanced PKA phosphorylation. Exendin 9-39 attenuated most responses, whereas insulin at comparable glycemic control reproduced few of these benefits, indicating effects that are at least partly GLP-1R dependent and not explained by glucose lowering.

We next compared our findings with previous studies of GLP-1 and the SAN. The key finding of Lubberding et al.^18^ was that, in pigs and isolated SAN preparations, GLP-1 enhanced calcium-clock-related phosphorylation through pacemaker-cell GLP-1R, shortened the pacing cycle, and accelerated diastolic depolarization; the effect persisted after autonomic or *I_f_* blockade, supporting a direct SAN action. However, that study used normal tissue and acute GLP-1 exposure and therefore could not determine whether chronic GLP-1RA treatment can reverse established structural injury. In STZ-SND rats, we found that eSuba still increased IHR after autonomic blockade, consistent with a direct action; moreover, chronic benefits were accompanied by reversal of fibrotic, inflammatory, oxidative, and glial programs, indicating that GLP-1R activation may act not only on the pacemaker-cell calcium clock but also on the diseased microenvironment. Anti-inflammatory IL-10 therapy also improves SAN function in STZ mice^29^, consistent with our reduction in CCR2a^+^/CD68^+^ infiltration; unlike a single-cytokine approach, however, eSuba also reversed oxidative, metabolic, and structural remodeling. Regarding the autonomic controversy^19, 20^, the persistence of increased IHR after autonomic blockade supports a direct component, whereas changes in PNS/SNS indices indicate that autonomic modulation also contributes to the overall heart-rate response; these mechanisms are not mutually exclusive and their relative weight may vary with species, dose, duration, and disease state.

CaMKII is an important regulator of basal SAN function. Previous studies have mainly shown that Met281/282 oxidation of CaMKII is a pathogenic step in SND^13–15^; whereas interventional evidence has largely come from inhibition or genetic editing of CaMKII oxidation^30^. The present study, from the perspective of therapeutic GLP-1RA intervention, observed a decrease in ox-CaMKII and its ratio to total CaMKII, suggesting that GLP-1R activation can indirectly reduce this oxidative modification. In the present study, eSuba significantly reduced ox-CaMKII and its ratio to total CaMKII while increasing SIRT3, a key node in mitochondrial antioxidant defense and deacetylation regulation; GLP-1RAs have been shown to exert cardioprotection through SIRT3-dependent pathways^31^. Importantly, we did not perform SIRT3 inhibition or knockdown, so the relationship between SIRT3 and ox-CaMKII should be regarded as associative and its causal role remains to be established. The role of CaMKII in diabetic cardiomyopathy and its post-translational regulation has been repeatedly emphasized^32, 33^. In addition, DRP1 expression in the eSuba group was lower than in the H_2_O_2_ model group (**Figure 4g,i**), suggesting reduced mitochondrial fission and a shift toward mitochondrial dynamic homeostasis; together with reports that GLP-1RAs improve myocardial metabolism and remodeling through NR4A1^34^, eSuba may improve both mitochondrial antioxidant capacity and dynamics in the SAN. However, mitochondrial morphology and function were not directly measured in this study and require further validation.

eSuba increased Ca_v_1.3, SERCA2a, and p-RyR2 (Ser2808) levels and increased the p-PKA/PKA ratio (**Figure 6c,d**); transcriptomics also showed upregulation of cAMP-synthesis genes and downregulation of phosphodiesterase genes (**Figure 6a,b**). Ser2808 is a PKA consensus phosphorylation site in RyR2, and increased phosphorylation at this site can enhance the calcium sensitivity of RyR2 and promote sarcoplasmic reticulum calcium release and recovery of calcium cycling. Therefore, an increased p-RyR2/RyR2 ratio more likely reflects PKA pathway activation and normalization of calcium cycling than a pathogenic phosphorylation event^35^. Previous studies have suggested that GLP-1R is expressed in SAN pacemaker cells and can directly regulate their function^17, 21^; our human SAN single-cell data and mouse SAN immunofluorescence further suggest relative enrichment of GLP-1R in the pacemaker cell population (**Figure S10**). Together with the changes in the cAMP-PKA gene set and p-PKA/PKA, we speculate that eSuba enhances RyR2 Ser2808 phosphorylation through the GLP-1R–cAMP–PKA axis; however, we did not directly measure cAMP or PKA activity, so this pathway is currently supportive evidence, and roles of other kinases or phosphatases cannot be excluded^16, 17, 36^. It should be noted that p-PKA/PKA was also increased in the insulin group, indicating that PKA phosphorylation is not specific to GLP-1R. However, the effects of eSuba on calcium-handling proteins and RyR2 phosphorylation were accompanied by reversal of multiple histological and functional indices by the GLP-1R antagonist, and the cAMP-PKA gene set showed directionally consistent changes at the transcriptomic level, supporting the PKA pathway as an important, though not the sole, component of eSuba action. Because Ca_v_1.3 is required for β-adrenergic acceleration of SAN firing^37^ and is functionally coupled to RyR2-dependent Ca^2+^ release^38^, the higher Ca_v_1.3 protein after eSuba may contribute to chronotropic reserve. Measuring Ca_v_1.3 current, surface expression, and β-adrenergic responsiveness would help establish this link and determine whether the protective effect of eSuba is mediated by Ca_v_1.3.

Maintenance of SAN function depends on electrical coupling between pacemaker cells and between pacemaker cells and atrial myocardium. Connexin distribution in the mouse SAN is closely related to conduction properties^39^; desmosomal junctions are indispensable for adult SAN function^40^; and intercalated disc/adherens junction genes such as MYZAP are linked to conduction abnormalities^41, 42^. In the present study, the SAN transcriptome in the eSuba group showed *Myzap* upregulation and favorable changes in junction-related genes such as *Klf2*/*Gja5* (**Figure 5a**), accompanied by restoration of Na_v_1.5, Ca_v_1.3, SERCA2a, and other electrical/calcium-handling proteins (**Figure 5c,f**), suggesting that eSuba not only improves the calcium clock and membrane clock of pacemaker cells but may also promote overall rhythm recovery by improving intercellular coupling and impulse output. Consistent with this, Cx43 density was reduced and its distribution shifted from intercalated discs to the lateral membrane in the SND group, both improved by eSuba (**Figure S9**). Together with *Myzap* upregulation in the transcriptome (**Figure 5a**), these findings suggest that eSuba may improve connexin expression and distribution, enhance intercellular coupling, and promote impulse output; however, direct evidence from Cx43 protein quantification and electrical conduction mapping is still needed.

The SAN is not composed solely of pacemaker cells but contains abundant neural, glial, and interstitial components, and its "brain-like" architecture participates in pacemaker regulation^2^. Glial cells of the intrinsic cardiac nervous system support neurotransmitter homeostasis, axonal integrity, and immune surveillance, and their abnormal activation is associated with arrhythmia and heart failure^43^; the SAN innervation atlas further reveals a complex neuro-glial network^44^. In this study, Schwann/glial-related transcripts such as *Mpz*, *Pmp22*, *Ncmap*, *Gjb1*, *S100b*, and *Kcna1* were increased in the SND SAN and decreased after eSuba treatment (**Figure 6e,f**), suggesting that pathological glial activation/myelin remodeling is a previously under-recognized component of SND that can be modified by GLP-1RA therapy. Given the broad expression of GLP-1R in neural tissue and its neuroprotective effects^19, 45^, the effects of eSuba on glial programs may reflect both improvement of the local inflammatory/oxidative environment and direct modulation of the neuro-glial unit. Because we did not perform protein validation of glial markers (such as S100β, GFAP, or MPZ) or neural electrophysiological recordings, this inference remains a hypothesis and should be tested in future studies using lineage tracing, glial-specific manipulation, and ex vivo neuro-SAN co-culture.

### Clinical implications

eSuba has been approved for type 2 diabetes and has a favorable pharmacokinetic and safety basis^25–28, 46^; its cardiovascular benefits and metabolic improvements have been further supported by recent clinical studies^27^. The present study suggests that GLP-1RAs may have therapeutic potential in metabolic stress-associated SND beyond glucose and weight reduction; in addition, the heart-rate-raising effect of GLP-1RAs may be therapeutically meaningful in patients with SND, contrasting with the traditional view that this effect is a potential risk in heart failure populations^47^. These inferences require validation in larger animal models and in humans.

### Limitations

Several limitations should be noted: (1) we did not perform patch-clamp recording, calcium imaging, or optical mapping in isolated SAN or primary pacemaker cells, so specific ionic currents and calcium-clock changes could not be directly determined; (2) the study was performed only in a rodent model and requires validation in human-like/large-animal models and in humans; (3) Body weight, food intake, and blood pressure were not measured. Weight loss and blood pressure reduction can alter heart rate and autonomic tone, and energy balance/obesity can affect systemic metabolic and inflammatory status; central autonomic modulation may further influence these responses^48^. These variables may therefore confound the heart-rate and remodeling endpoints; and (4) bulk RNA-seq cannot distinguish changes in cell composition from changes in intracellular expression. These limitations do not affect the main conclusions of this study but indicate that the underlying mechanisms require further investigation.

## Conclusions

The long-acting GLP-1RA eSuba improves STZ-induced SAN dysfunction by attenuating electrophysiological and structural remodeling: pacemaker function improved in both the acute and chronic phases, together with reversal of multiple transcriptional programs, reduced local fibrosis and inflammation, decreased ox-CaMKII, increased SIRT3, and restoration of calcium-handling proteins and PKA phosphorylation. Exendin 9-39 attenuated most responses, supporting at least partial GLP-1R mediation, and the insulin active comparator, despite comparable glycemic control, did not reproduce most of these benefits, indicating that they are not explained by glucose lowering. These results provide an experimental basis for pharmacological intervention in metabolic stress-associated SND and suggest that the associated molecular changes—ox-CaMKII, SIRT3, PKA/calcium handling, and glial programs—warrant further mechanistic and translational investigation.

## Acknowledgments

H.F.L., J.W., and B.L. conceptualized the study; H.F.L. developed the methodology; H.F.L., Y.Z., R.Z., W.F., C.L., and Z.L. performed the investigation; H.F.L., Y.Z., R.Z., and W.F. performed the formal analysis; H.F.L., Y.Z., and R.Z. curated the data; H.F.L., Y.Z., and R.Z. wrote the original draft; H.F.L., Y.Z., J.W., and B.L. reviewed and edited the article; H.F.L. created the visualization; J.W. and B.L. supervised the research; J.W., B.L., and H.F.L. acquired the funding. All authors approved the final version.

## Data Availability

The RNA-seq data generated in this study are available from the corresponding author upon reasonable request.

## Supplemental Material

Supplemental Methods; **Figures S1–S15** (with legends).

